# The tubulin repertoire of *C. elegans* sensory neurons and its context dependent role in process outgrowth

**DOI:** 10.1101/075879

**Authors:** Dean Lockhead, Erich M. Schwarz, Robert O'Hagan, Sebastian Bellotti, Michael Krieg, Maureen Barr, Alexander R. Dunn, Paul W. Sternberg, Miriam B Goodman

## Abstract

Microtubules contribute to many cellular processes, including transport, signaling, and chromosome separation during cell division (Kapitein and Hoogenraad, 2015). They are comprised of αβ-tubulin heterodimers arranged into linear protofilaments and assembled into tubes. Eukaryotes express multiple tubulin isoforms (Gogonea *et al.*, 1999), and there has been a longstanding debate as to whether the isoforms are redundant or perform specialized roles as part of a tubulin code (Fulton and Simpson, 1976). Here, we use the well-characterized touch receptor neurons (TRNs) of *Caenorhabditis elegans* to investigate this question, through genetic dissection of process outgrowth both *in vivo* and *in vitro*. With single-cell RNA-seq, we compare transcription profiles for TRNs with those of two other sensory neurons, and present evidence that each sensory neuron expresses a distinct palette of tubulin genes. In the TRNs, we analyze process outgrowth and show that four tubulins (*tba-1*, *tba-2*, *tbb-1*, and *tbb-2*) function partially or fully redundantly, while two others (*mec-7* and *mec-12*) perform specialized, context-dependent roles. Our findings support a model in which sensory neurons express overlapping subsets of tubulin genes whose functional redundancy varies between cell types and *in vivo* and *in vitro* contexts.

**Highlight Summary:** Microtubules contribute to key cellular processes and are composed of αβ-tubulin heterodimers. Neurons in *C. elegans* express cell type-specific isoforms in addition to a shared repertoire and rely on tubulins for neurite outgrowth. Isoform function varies between *in vivo* and *in vitro* contexts.

**Abbreviations:**

TRNs: Touch Receptor Neurons
RNA-seq: RNA sequencing
RFG: Receptive Field Gap
TPM: Transcripts per Million
ECM: Extracellular Matri**x,**
CV: Coefficient of Variation

**Conflict of Interest:** The authors declare no conflicting financial interests.

## Introduction

Shortly before the discovery that microtubules (MTs) are composed of tubulin subunits (Mohri, 1968), investigators found differences in the chemical properties and thermal stability of various microtubules (Behnke and Forer, 1967). These experiments, along with experiments showing the inducible expression of chemically-distinct tubulins (Fulton and Simpson, 1976), led to the formulation of the multi-tubulin hypothesis which posits that there are multiple tubulin isoforms within cells and that each one has a distinct function (Fulton and Simpson, 1976).

Early experiments in the budding yeast *Saccharomyces cerevisiae* argued against the multi-tubulin hypothesis by showing that its two α—tubulins can function redundantly (Schatz *et al.*, 1986). Similarly, the β—tubulins of *Aspergillus nidulans* were also found to be interchangeable (May, 1989). In contrast, Hoyle and Raff observed isoform-dependent function in *Drosophila* (Hoyle and Raff, 1990). In particular, they demonstrated that the function of the sole β-tubulin isoform expressed in the male testes, β2, cannot be rescued by expression of the β3 isoform and that the β3 isoform acts in a dominant-negative fashion when co-expressed with β2 at levels exceeding 20% of the total tubulin.

The observation that most studies in isolated cells demonstrated functional redundancy whereas the majority of studies in multicellular organisms showed isoform-specific function led to the question of whether isoform-specific function may be context-dependent (Ludueña, 1993; Hurd *et al.*, 2010). In this study, we leverage the genetically-tractable model organism *C. elegans* and its well characterized and compact nervous system to address this question with respect to the role that MTs play in axon outgrowth. We use RNA-seq to survey the genes expressed in single, identified mechanoreceptor, thermoreceptor, and chemoreceptor neurons and mine the resulting datasets to determine tubulin genes expressed in each type of neuron. Leveraging this information, we manipulate the tubulin composition of the well-characterized touch receptor neurons (TRNs), both *in vivo* and *in vitro,* and examine the consequences of these alterations on the outgrowth of their neural processes.

*C. elegans* nematodes rely on six TRNs to sense gentle touch along the length of their bodies (Chalfie and Thomson, 1979; Chalfie *et al.*, 1985). The lateral TRNs, ALML/R and PLML/R, arise embryonically, whereas the ventral neurons, AVM and PVM, arise later in larval development (Chalfie and Sulston, 1981). During development, the TRN cell bodies migrate towards their adult positions and then extend processes anteriorly, and in the case of the PLMs, posteriorly as well (Chalfie and Sulston, 1981). The anteriorly-directed PLM process grows rapidly upon hatching and then pauses in a SAX-1/SAX-2-dependent manner before resuming growth (Gallegos and Bargmann, 2004). This pause has been shown to be crucial for the proper tiling of the body surface (Gallegos and Bargmann, 2004).

TRN axons are packed with a distinctive bundle of 15-protofilament microtubules that depend on the expression of MEC-12 α-tubulin and MEC-7 β-tubulin (Chalfie and Thomson, 1979, 1982; Chalfie and Au, 1989; Savage *et al.*, 1989). These unusual microtubules are required for gentle touch sensation (Chalfie and Sulston, 1981) and to regulate gene expression (Bounoutas *et al.*, 2011), but not for the generation of electrical currents in the TRNs in response to a mechanical stimulus (O’Hagan *et al.*, 2005; Bounoutas *et al.*, 2009). MEC-12 α-tubulin is the only *C. elegans* tubulin subject to acetylation at lysine 40; this modification is needed for normal touch sensation (Akella *et al.*, 2010; Shida *et al.*, 2010; Topalidou *et al.*, 2012) and to constrain the number of protofilaments in each microtubule to fifteen (Cueva *et al.*, 2012).

Unlike *mec-7* and *mec-12*, the expression profile of other tubulin isoforms in TRNs and their contribution to the development and function of these cells is poorly understood. Of the other eight α-and five β-tubulins in the *C. elegans* genome (Consortium, 1998; Gogonea *et al.*, 1999), previous investigators have found evidence for the expression of two other α-tubulins in the TRNs in addition to MEC-12: *tba-1* and *tba-2* (Fukushige *et al.*, 1993, 1995).

Coupled with visualization of tubulin gene expression *via* transcriptional GFP fusions and CRISPR/Cas9-mediated insertion of TagRFP-T into the genome, our study indicates that each neuron expresses a distinct palette of tubulin isoforms. The TRN mechanoreceptors express transcripts encoding three α-and three β-tubulins in addition to *mec-7* and *mec-12.* We analyze the effect of null mutations in a subset of these isotypes *in vivo* and *in vitro* and demonstrate that *mec-7* is required for the wild type outgrowth of TRN processes *in vivo* and *in vitro*, while *mec-12* plays a partially redundant role with *tba-1* in *in vivo* and is required for wild type outgrowth in *in vitro*.

## Results

To study the effects of tubulin isoforms on TRN process outgrowth, we first sought to understand the contribution of microtubules to the outgrowth of TRNs *in vivo*. Previous work established that growing *C. elegans* on media containing the microtubule-destabilizing drug, colchicine, was sufficient to eliminate microtubules in the lateral TRNs (Chalfie and Thomson, 1982; Bounoutas *et al.*, 2011). We thus examined the effect of colchicine on TRNs of worms expressing one of two TRN-specific GFP transgenes, *uIs30* or *uIs31* (O’Hagan *et al.*, 2005; Bounoutas *et al.*, 2009). As expected from prior work (Chalfie and Thomson, 1982), colchicine impaired touch sensation: untreated animals responded to an average of 8.2 ± 0.16 (mean ± SEM, *N =* 75 animals) of 10 trials, while colchicine-treated animals responded to 2.3 ± 0.17 of 10 trials (mean ± SEM, *N* = 75 animals).

Within each cohort of animals, the position (normalized to body length) of the ALM and PLM cell bodies and the endpoints of their processes were highly stereotyped (Figure 1). The average lengths of the anterior processes of ALM and PLM were similar to those reported previously in untreated, wild-type animals (Gallegos and Bargmann, 2004; Shida *et al.*, 2010; Petzold *et al.*, 2013): 382 ± 12 µm (mean ± SEM, *n* = 20) and 498 ± 18 µm (mean ± SEM, *n* = 20) for ALM and PLM, respectively. Colchicine treatment shifted PLM cell bodies posteriorly and shortened both the anterior and posterior PLM processes (Figure 1A, 1C, 1E). The defect in PLM positioning and outgrowth resulted in an increase in the receptive field gap between the ALM and PLM neurons (Figure 1A, B). Colchicine treatment also displaced the ALM cell body posteriorly (Figure 1C), but did not appear to affect the length of the anteriorly directed ALM process (Figure 1D). The origin of this positioning defect is unclear. ALM, but not PLM cell bodies migrate a considerable distance posteriorly during development (Sulston and Horvitz, 1977; Manser and Wood, 1990), suggesting that colchicine treatment may affect posteriorly-directed cell migrations. Taken together with prior work establishing that colchicine-treated animals lack 15-protofilament microtubules in both ALM and PLM (Chalfie and Thomson, 1982), these results show that disrupting MTs has surprisingly mild effects on the extension of TRN processes *in vivo*.

**Figure 1.**
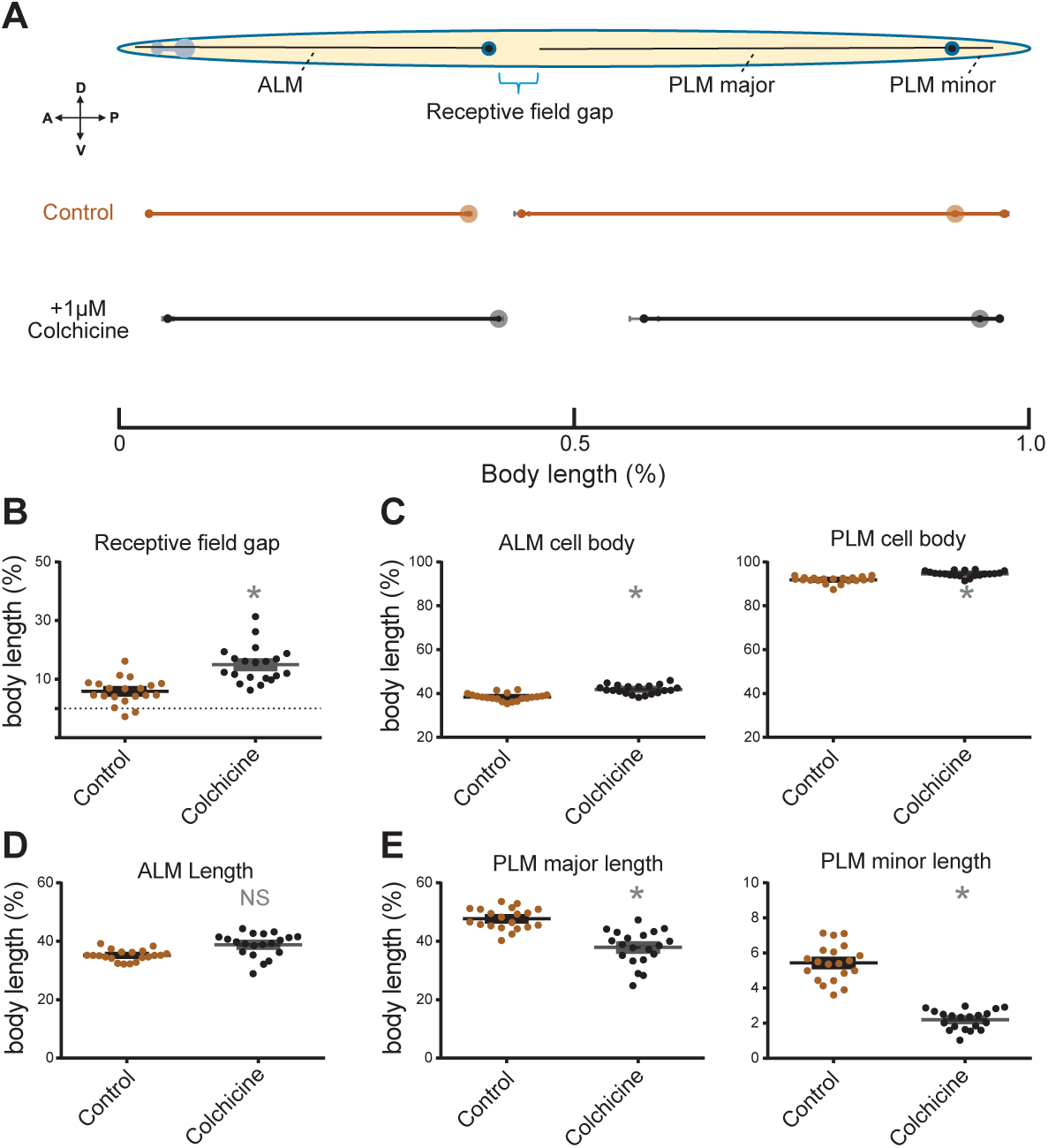
Pharmacological disruption of microtubules disrupts PLM length *in vivo*. **A.** Normalized results from average of 20 control *uIs31[mec-17p::gfp]* and colchicine-treated animals grown from embryos for three days. Body length measured from mouth to tail. Dark circles on schematic indicate cell bodies. Anterior is left, ventral is down. **B-E.** Growth and cell position parameters from all 20 animals in each treatment group expressed as a fraction of the body length. *: indicates *p* < 0.05 as determined by t-test for a given parameter between control and colchicine-treated groups.

### TRN gene expression profiling

Having established that microtubules contribute to the outgrowth of the posterior TRNs *in vivo*, we sought to identify the tubulin genes that are expressed in these TRNs. To reach this goal, we used single-cell RNA sequencing (RNA-seq) to generate gene expression profiles of PLM neurons harvested from late L4 larvae. We took two steps to differentiate between tubulin genes expressed specifically in the TRNs and those expressed more broadly. First, we compared expression levels to a dataset reported previously for mixed-stage *C. elegans* larvae (Schwarz *et al.*, 2012). Second, we generated gene expression profiles for two other neurons: a chemosensory neuron, ASER, and a thermosensory neuron, AFD. We detected expression of 5,086 protein-coding genes in PLM neurons (Figure 2A, Tables S3, S4) and similar numbers in AFD and ASER neurons (5,824 and 4,601 genes, respectively) isolated from animals late in larval development or in early adulthood (Figure 2, Tables S3, S4). These results are similar to those reported for other neurons isolated from larvae and analyzed by RNA-seq (Spencer *et al.*, 2014). Thus, on average, individual classes of *C. elegans* neurons appear to express approximately 5,000 genes or one-quarter of the genome.

**Figure 2.**
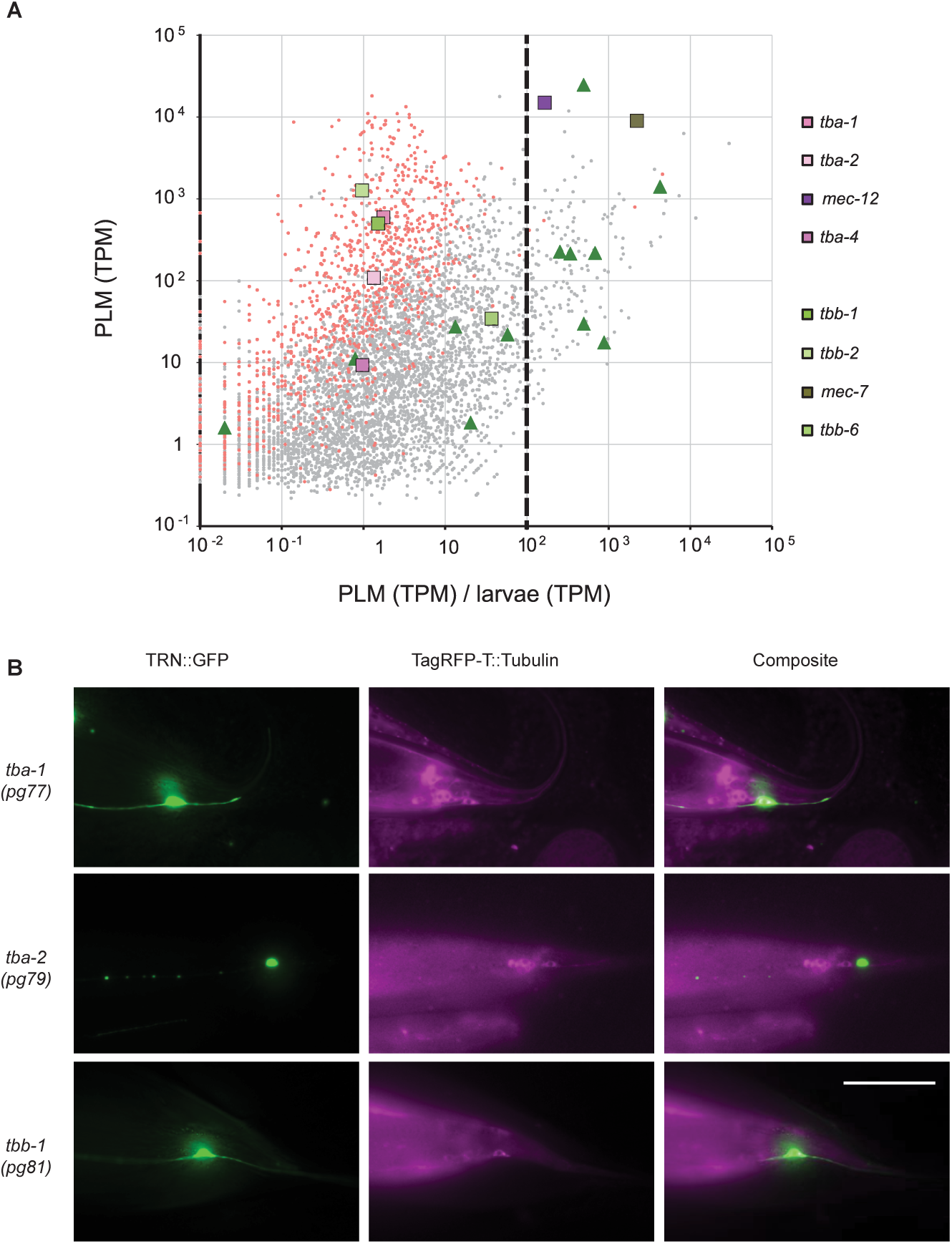
Tubulin isoform expression in PLM. **A.** Expression of 5,099 *C. elegans* genes (5,086 protein-coding and 13 ncRNA-coding) that exhibited above-background expression in PLM (Methods and Supplementary Table S4). Gene expression, measured in transcripts per million (TPM), is plotted against expression in PLM relative to expression in whole larvae. Each gray or red dot represents a single gene expressed in PLM; red dots are 1,060 housekeeping genes (Schwarz *et al.*, 2012). Genes encoding 8 tubulins are highlighted as larger squares and those encoding 12 non-tubulin *mec* genes are shown as larger green triangles. The dashed vertical line highlights 100-fold more transcripts in PLM than in whole larvae. **B**. Co-expression of α-tubulin and β-tubulin TagRFP-T protein fusions and *uIs31* TRN::GFP transcriptional fusion. Left: *uIs31(mec-17p::gfp)* showing PLM. Center: TagRFP-T tubulin fusion protein. Right: Composite image. TagRFP-T was false colored as magenta for enhanced visibility. Expression of isoforms in TRNs seen as white in composite overlap of *uIs31* marker and TagRFP-T::Tubulin. Images are oriented so that the anterior of the animal is on the left and ventral side is on the bottom. Scale bar: 50 µm.

The number of genes expressed in these three sensory neurons was roughly half that detected in mixed-stage whole *C. elegans* larvae, which we used as a control for general, non-PLM-specific gene activity (9,918 genes). Of the protein-coding genes expressed in PLM, 1,060 (21%) were housekeeping genes by previously defined criteria (Schwarz *et al.*, 2012). Conversely, 955 genes (19%) could be defined as PLM-enriched, 1,089 (24%) as ASER-enriched, and 1,203 (21%) as AFD-enriched, by the criterion that each gene exhibited at least 10-fold more expression in the relevant sensory neuron than in whole larvae (Table S4). By these criteria, PLM-enriched genes included these genes that mutate to disrupt touch sensation (in decreasing order of expression ratios): *mec-18*, *mec-7, mec-10*, *mec-1*, *mec-4*, *mec-17*, *mec-2*, *mec-9*, *mec-12, mec-6*, *mec-14* and *mec-8*. AFD-enriched genes included *gcy-8*, *gcy-23*, *gcy-18*, *hen-1*, and *ttx-1,* while ASER-enriched genes included *gcy-22, gcy-19*, and *flp-6*. Consistent with its long cilium, transcripts encoding subunits of the BBSome, IFT-A, and IFT-B ciliary transport complexes were enriched in ASER by an average of 27-, 88-and 100-fold, respectively. A similar enrichment was not detected in either AFD (which has a very short cilium) or in PLM, which lacks ciliated structures entirely. Thus, the application of RNA-seq technology to single, identified neurons extracted from older nematodes is sufficient to detect neuron-specific genes.

*C. elegans* has nine α-and six β-tubulin genes in its genome (Consortium, 1998; Gogonea *et al.*, 1999). We detected transcripts for eight of these fifteen tubulins in PLM neurons. In decreasing order of PLM-specific expression, they were *mec-7*, *mec-12*, *tbb-6*, *tba-1*, *tbb-1*, *tba-2*, *tba-4*, and *tbb-2* (Figure 2A). Because we detected only low levels of expression for *tba-4* and *tbb-6* in the PLMs (9 and 34 transcripts per million [TPM], respectively), we focused on the other tubulin isoforms in this study. In the AFD neuron, we detected nine isoforms (*tba-1, tba-2, mec-12, tba-4, tbb-1, tbb-2, mec-7, tbb-4,* and *tbb-6*; Figure S1) and in the ASER neuron, we detected eight (*tba-1, tba-2, mec-12, tba-4, tbb-1, tbb-2, mec-7,* and *tbb-4*; Figure S1). These results are consistent with a previous study that found *tbb-4* expression in AFD and ASE (Hurd *et al.*, 2010); however, in contrast with their results, we did not observe expression of *tba-9* in either ciliated neuron. In AFD and ASER, we also detected significant expression of the γ**-**tubulin, *tbg-1* (Figure S1, Table S4). Based on this analysis, the PLM mechanoreceptors, ASER chemoreceptors, and AFD thermoreceptors all appear to co-express transcripts encoding three α-tubulins (*tba-1, tba-2*, *mec-12*) and three β-tubulins (*tbb-1*, *tbb-2, mec-7*).

The PLM neurons expressed at least 10-fold more transcripts encoding all tubulins than either AFD or ASER (Figure 2A, S1, Table S4). This difference in total tubulin transcript expression is primarily due to the large increase in *mec-7* and *mec-12* transcript numbers observed in PLM compared to AFD and ASER. We detected ~15,000 TPM and ~9,000 TPM for *mec-12* and *mec-*7, respectively, in the PLM (Table S4). In contrast, the highest-expressed isoform in either AFD or ASER is *tbb-2* in AFD at 889 TPM, roughly an order of magnitude lower. After *mec-7*, *tbb-2* was also the most highly expressed β-tubulin in the PLM (Figure 2A, S1). Similarly, *tba-1* was the most highly expressed α-tubulin in AFD and ASER, as well as in PLM, after *mec-12*. We found expression of *tbb-4* in both AFD and ASER, but not in PLM (Figure S1). *tbb-4* has been previously shown to be expressed in ciliated neurons and to be required for their morphology and for behaviors that depend on these neurons (Portman and Emmons, 2004; Hurd *et al.*, 2010; Hao *et al.*, 2011). Collectively, these data suggest that each sensory neuron has a distinct tubulin expression palette, and that the 10-to 100-fold amplification in the expression of *mec-12* and *mec-7* in the TRNs is consistent with the central role that these tubulins play in TRN development and function.

### Tubulin genes expressed in TRNs

Next, we sought to characterize the cells and tissues expressing the tubulin isoforms detected in TRNs. Because the expression patterns of *mec-7* and *mec-12* are well established (Chalfie and Sulston, 1981; Chalfie and Thomson, 1982; Savage *et al.*, 1989, 1994; Hamelin *et al.*, 1992; Fukushige *et al.*, 1999; Bounoutas *et al.*, 2009), we did not examine them further. Instead, we focused on *tba-1*, *tba-2*, *tbb-1*, and *tbb-2,* and used CRISPR/Cas9-mediated gene editing to insert a fluorescent protein into the native loci as well as conventional transcriptional fusions to visualize their expression.

Using a self-excising cassette strategy (Dickinson *et al.*, 2013), we inserted sequences encoding TagRFP-T into the *tba-1, tba-2,* and *tbb-1* loci, creating the *pg77*, *pg79*, and *pg81* insertion alleles expressing TagRFP-T fusion proteins. (Efforts to insert the same tag into the *tbb-2* locus were unsuccessful.) In lines tagging the amino terminus of TBA-1, TBA-2 and TBB-1, we detected fluorescence in body wall muscle, pharynx, intestine, and in neurons in the head, tail, and ventral nerve cord. Throughout the body, TagRFP-T fluorescence was considerably higher for *tba-1* and *tbb-1* insertions than they were for *tba-2*. These variations in apparent expression levels could reflect our finding that *tba-2* transcript counts were lower than those for *tba-1* and *tbb-1* in all three of the neurons examined with RNA-seq (Figure 2A, Figure S4).

To determine whether or not these tubulins were expressed in the TRNs, we combined the *pg77*, *pg79*, and *pg81* TagRFP-T insertion alleles with *uIs31* that drives expression of GFP exclusively in the TRNs. We found TBA-1 and TBB-1 in all six TRNs, but could not detect TBA-2 in any TRNs with this method (Figure 2B). We also generated conventional transcriptional fusions and obtained lines expressing GFP downstream of the putative promoter sequences for *tba-1*, *tba-2*, and *tbb-2*. (Efforts to visualize expression from the *tbb-1* promoter were unsuccessful.) We detected fluorescence from expression of all three promoters throughout the body (Figure S2), as well as in the TRNs (Figure S3), including from the *tba-2* promoter.

In summary, we tested for TRN expression using two complementary, fluorescent protein-based visualization methods. For *tba-1*, fluorescence was visible both from an insertion allele and from expression of a transcriptional fusion. For *tba-2*, fluorescence was not visible from the insertion allele, but it was visible from expression of a transcriptional fusion. Even though only one visualization method was successful for *tba-2, tbb-1* and *tbb-2*, we suggest that these results indicate that the TRNs express *tba-1, tba-2, tbb-1*, and *tbb-2*.

### *mec-7* is necessary for wild type TRN outgrowth *in vivo*

To study the contribution of different tubulin isoforms to TRN outgrowth, we created animals with putative null mutations for one or more tubulin isoforms in the presence of either *uIs31* or *uIs30* transgenes that drive GFP expression with the TRN-specific promoter of the *mec-17* gene. Figure 3 compares the positions of the ALM and PLM cell bodies and their process terminations in control animals and in single α-tubulin mutants (*tba-1, tba-2,* and *mec-12*), single β-tubulin mutants (*tbb-1*, *tbb-2*, and *mec-7*), and selected double α-tubulin and β-tubulin mutants. We were unable to examine *tba-1tba-2, tba-1;tbb-1, tba-1;tbb-2,* or *tbb-2tbb-1* double mutants due to their embryonic lethality (Phillips *et al.*, 2004; Baran *et al.*, 2010).

**Figure 3.**
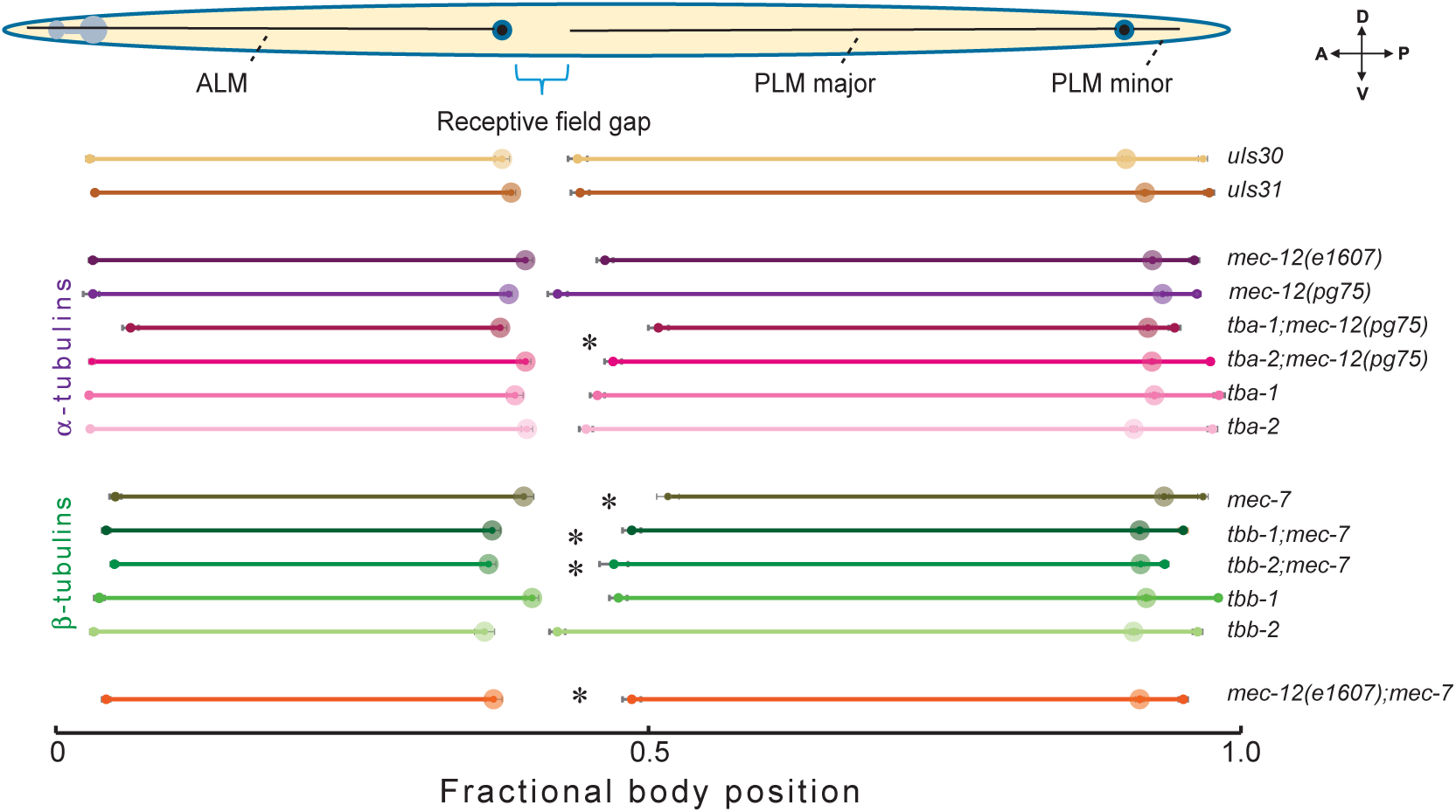
Mutation of *mec-7* or combination of *mec-12* with *tba-1* or *tba-2* mutations increases the receptive field gap. Normalized results from average of 20 animals imaged for each genotype as indicated. Anterior left, ventral down. *Indicates a statistically significant increase in RFG compared to respective GFP control (*p* < 0.05). Table 1 lists the values for all of these parameters.

Of these six tubulin genes, *mec-7* had the strongest effects on TRN process outgrowth (Figure 3, Table 1). Loss of *mec-7* nearly tripled the length of the receptive field gap (from 4.2 ± 1% to 12.2 ± 0.8% body length), an effect that arises primarily from defects in the positioning of the PLM cell body and its process outgrowth. Animals carrying the *e1607* and *pg75* null alleles of *mec-12* had slightly longer PLM process than wild type, but those carrying null mutations in *tba-1, tba-2, tbb-1,* or *tbb-2* had essentially wild-type processes (Figure 3, Table 1). Notably, *tba-1;mec-12(pg75)* double mutants had a larger receptive field gap (13.3±0.9% body length) than either single mutant and were the only example of genetic enhancement among the double mutants we examined.

**Table 1.** Position of the ALM and PLM cell bodies and their process lengths *in vivo* as a function of genotype. *uIs31* and *uIs30* are wild-type, control animals expressing GFP exclusively in the TRNs. Data are mean±sem (*n=*20) and reported as a percentage of body length, except for body length. One-way ANOVA was used to determine the effect of genotype and applied separately to strains containing either the *uIs31* or *uIs30* transgenes, followed by Dunnett’s multiple comparisons test between individual genotypes and their GFP control. Values in bold are those that differed from control at the *p <* 0.05 level.

| Genotype | Body length ($\mu\text{m}$) | Cell body position | | Process length (%) | | | Receptive field gap (%) |
| --- | --- | --- | --- | --- | --- | --- | --- |
|  |  | ALM | PLM | ALM | PLM (Major) | PLM (Minor) |  |
| *uIs31[pmec-17::gfp]* | 1148±24 | 38.4±0.4 | 91.9±0.3 | 35.1±0.4 | 47.7±0.8 | 5.4±0.2 | 5.8±1 |
| *uIs30[pmec-17::gfp]* | 1127±31 | 37.7±0.6 | 90.3±0.3 | 34.8±0.5 | 46.3±0.7 | 6.5±0.3 | 6.4±0.9 |
| *uIs30;*<br>*mec-12(e1607)* | 1149±27 | 38.2±0.5 | **93.4±0.4** | 35.1±0.6 | **51.1±0.9** | **2.9±0.2** | 4.1±1 |
| *mec-12(pg75)uIs31* | 1096±15 | 39.6±0.6 | 92.5±0.5 | 36.5±0.6 | 46.2±0.6 | **4.0±0.2** | 6.7±0.9 |
| *tba-1(ok1135); uIs31* | 1186±20 | 38.7±0.7 | 92.7±0.4 | 35.9±0.7 | 47.0±0.6 | 5.5±0.3 | 6.9±0.9 |
| *tba-1(ok1135);*<br>*mec-12(pg75)uIs31* | 895±15 | **37.5±0.6** | 92.1±0.4 | **31.7±1** | **41.3±0.8** | **2.2±0.2** | **13.3±0.9** |
| *tba-2(tm6948);*<br>*uIs31* | 1128±25 | 39.7±0.5 | 91.0±0.3 | 36.9±0.4 | 46.2±0.6 | **6.6±0.4** | 5±0.7 |
| *tba-2(tm6948);*<br>*mec-12(pg75)uIs31* | 1200±23 | 39.6±0.5 | 92.5±0.3 | 36.6±0.5 | 45.5±0.6 | 4.9±0.3 | **7.4±0.7** |
| *uIs31; mec-7(u142)* | 1145±33 | 39.5±0.8 | **93.5±0.4** | 34.5±0.7 | **41.9±0.8** | **3.3±0.2** | **12.2±0.8** |
| *uIs30; tbb-1(gk207)* | 1100±21 | **40.2±0.5** | **92.0±0.3** | 36.7±0.7 | 44.9±0.7 | 6.1±0.3 | 7.3±0.8 |
| *uIs30; tbb-1(gk207);*<br>*mec-7(u142)* | 1027±32 | 36.8±0.8 | **91.8±0.4** | **32.1±0.7** | **41.3±0.9** | **3.1±0.4** | **13.6±1** |
| *uIs30;tbb-2(gk130)* | 1148±20 | 36.2±0.8 | 90.9±0.3 | 33±0.7 | 48.6±0.6 | **5.4±0.3** | 6.1±1 |
| *uIs30;tbb-2(gk130);*<br>*mec-7(u142)* | 1234±24 | 36.5±0.6 | 91.5±0.3 | **31.2±0.8** | 44.5±1 | **2±0.2** | **10.6±1.3** |
| *uIs30;*<br>*mec-12(e1607);*<br>*mec-7(u142)* | 1053±26 | 36.9±0.7 | 91.5±0.3 | 32.7±0.7 | **42.9±0.7** | **3.7±0.3** | **11.6±0.8** |
| One-way ANOVA,<br>*uIs31* strains | *F*(6,133)<br>*P* | 2.455<br>0.0277 | 4.739<br>0.0002 | 7.620<br><0.0001 | 12.20<br><0.0001 | 28.52<br><0.0001 | 17.77<br><0.0001 |
| One-way ANOVA<br>for *uIs30* strains | *F*(6,133)<br>*P* | 4.166<br>0.0007 | 7.105<br><0.0001 | 8.185<br><0.0001 | 17.06<br><0.0001 | 35.31<br><0.0001 | 12.31<br><0.0001 |

The minor defects in TRN process growth found in *mec-12* null mutants and in *tba-1;mec-12* double mutants are surprising for at least two reasons. First, 95% of all α-tubulin transcripts detected in PLM encode *mec-12* and 99% of these transcripts encode either *mec-12* or *tba-1* (Table S4). Second, *mec-12* loss-of-function mutants are expected to lack the distinctive bundle of 15-protofilament microtubules present in wild-type TRNs (Chalfie and Au, 1989; Cueva *et al.*, 2012). Collectively, these observations suggest that other tubulins may compensate for the loss of *mec-12* and *tba-1* and that 15-protofilament microtubules are not essential for TRN outgrowth *in vivo*.

We used a ten-trial, classical gentle touch assay (Chalfie *et al.*, 2014) to assess TRN function in single and double tubulin mutants. Consistent with earlier work (Chalfie and Sulston, 1981), we found that null mutations in *mec-7* and *mec-12*, alone or in combination, caused severe defects in touch sensation (Figure 4). In contrast, touch sensation was indistinguishable from wild type in null mutants of the four other tubulins, and indistinguishable from the defects seen for *mec-12* or *mec-7* in double mutants. These results are consistent with an important role for *mec-12* and *mec-7* tubulins in TRN function.

**Figure 4.**
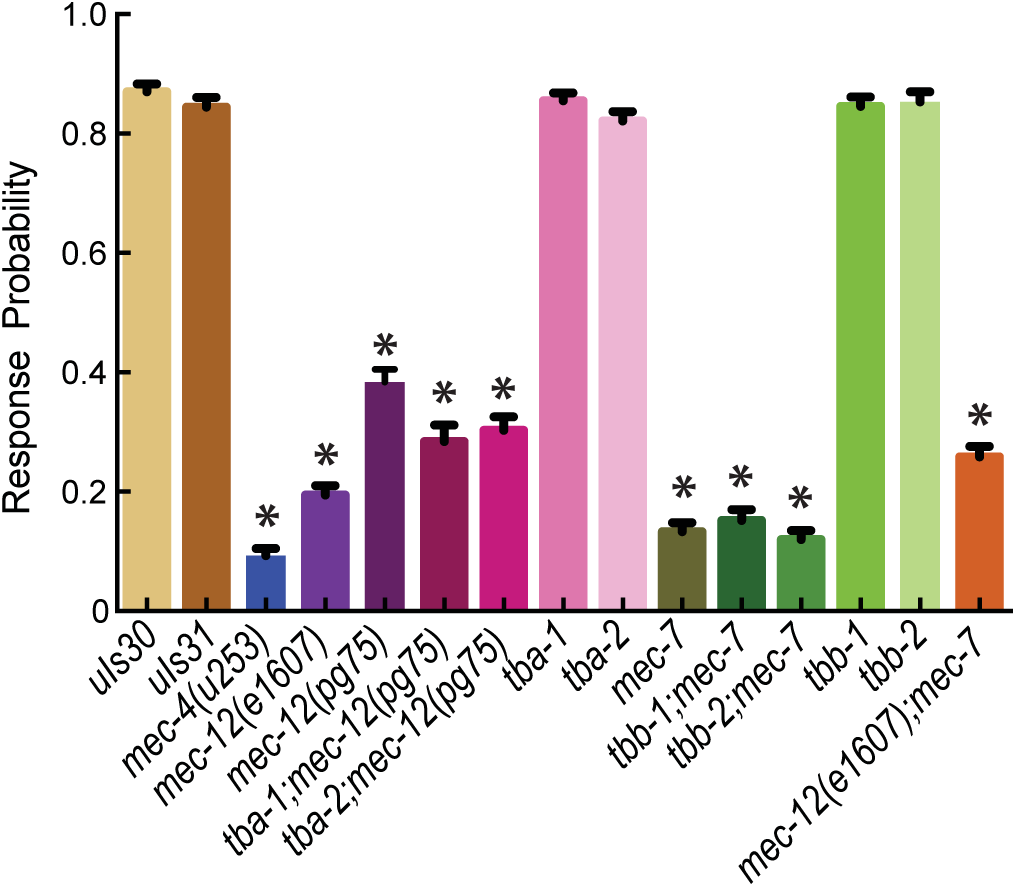
*tba-1, tba-2, tbb-1,* or *tbb-2* loss-of-function mutants have wild-type touch sensitivity. Results from gentle touch assays on 75 animals per genotype conducted blind to genotype on three separate days. Experimenter blinded to genotype. Statistical significance determined by omnibus ANOVA with respective GFP control, followed by Dunnett’s multiple comparisons test between individual genotypes and their GFP control. *indicates *p <* 0.05.

### Wild-type TRNs extend unipolar and bipolar processes *in vitro*

We sought to address the potential contribution of the ECM to process outgrowth and to explore the interplay between tubulin isoforms, the ECM, and process outgrowth by isolating the TRNs from their native context. To reach this goal, we first dissociated *uIs31* and *uIs30* control embryos in which GFP is expressed only in the TRNs (Methods) and characterized processes extended *in vitro*. TRN morphologies observed *in vitro* were broadly similar to those observed *in vivo* (Figure 5A) and to what has been reported previously (Zhang *et al.*, 2002; Strange *et al.*, 2007). The GFP-labeled cells fell into two categories, unipolar and bipolar, and extended three types of processes, unipolar, major bipolar, and minor bipolar (Figure 5A). We plotted the pooled data as cumulative probability distributions when making comparisons between genotypes, as seen in Figure 6B. The geometric means for control TRNs were 38.7 µm for the unipolar processes, 41.1 µm for the bipolar major processes, and 17.5 µm for the bipolar minor processes. We observed no effect of plating density on process length in the range of 0.5-1.5•10^5^ cells/dish (data not shown). On average, all three types of processes were approximately 10-fold shorter *in vitro* than *in vivo* (Figures 3,7, and Table 1).

**Figure 5.**
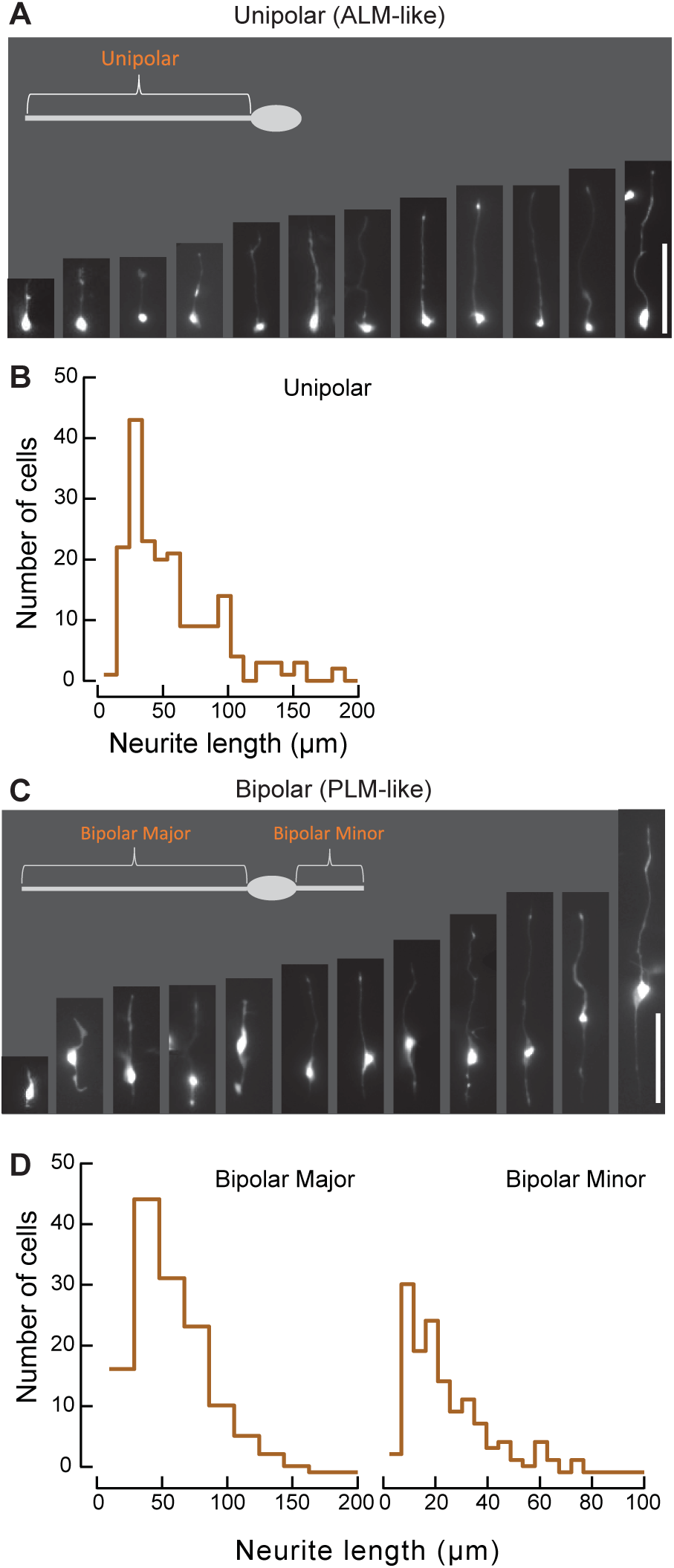
TRNs adopt similar morphologies *in vitro* as they do *in vivo*. **A.** Example images of unipolar and bipolar cells in culture. Processes are described as unipolar for single process on unipolar cells, as bipolar major for the large process on bipolar cells, and as bipolar minor for the small process. Scale bars: 50 µm. **B.** Histograms of process lengths for unipolar, major bipolar, and minor bipolar cells. Bin sizes were determined as in Shimazaki and Shinomoto (Shimazaki and Shinomoto, 2007).

**Figure 6.**
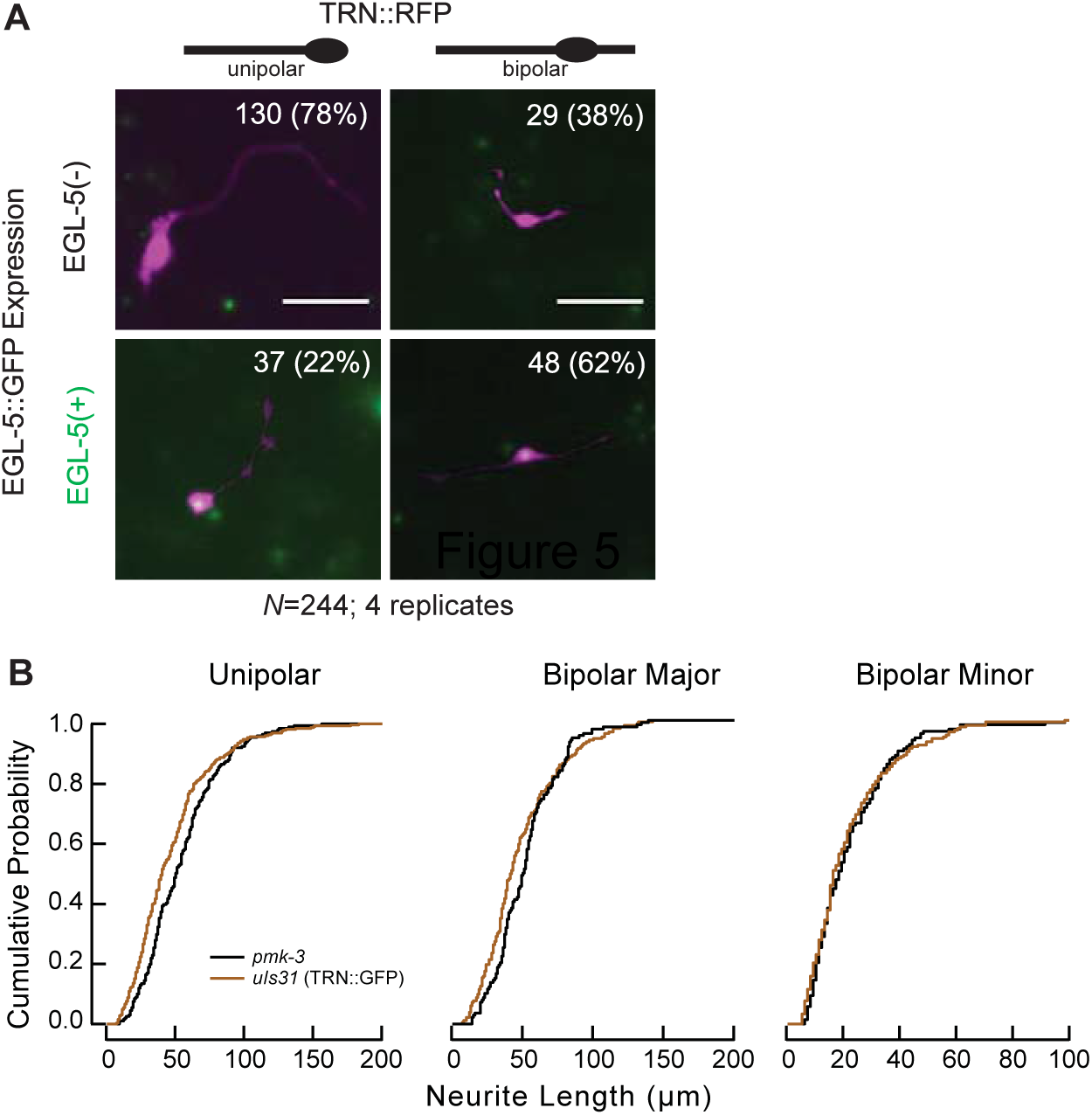
In *in vitro* culture, TRNs primarily adopt ALM-like cell fate and extend processes by a *pmk-3*-independent mechanism. **A.** Representative images of TRNs cultured from animals expressing TRN::RFP and EGL-5::GFP. RFP is false-colored as magenta, and GFP as green. Vertical columns from left to right: unipolar cells, bipolar cells. Horizontal rows from top to bottom: EGL-5(-), EGL-5(+) cells. Morphology and EGL-5 expression were scored manually. Scale bars: 50 µm. **B.** Cumulative probability distributions for TRNs isolated from control *uIs31[pmec-17::gfp]* and *pmk-3(ok169)* animals. Statistical significances were determined by Kolmogorov-Smirnoff tests.

Whereas *in vivo* process lengths have a small variance between individual animals and are normally distributed (Figure 1B-E), processes grown *in vitro* exhibit substantially larger variance in their length and their distribution is log-normal (Figure 5B). Indeed, control neuron lengths had coefficients of variation (CV) that were 6% and 80% for the anterior process of ALM *in vivo* and unipolar processes *in vitro*, respectively. Similarly, CV values were 8% for the anterior process of PLM *in vivo* and 90% for the major process of bipolar, PLM-like processes *in vitro*. This finding suggests that process length is tightly controlled *in vivo*, but not *in vitro*.

While TRN shapes were similar *in vitro* to those found *in vivo* where the ALM neurons are unipolar and the PLM neurons are bipolar, we sought to determine how *in vitro* morphology was connected to the cell fates of ALM and PLM. To address this question, we obtained *uIs115;uIs116* transgenic animals expressing RFP selectively in the TRNs and a GFP-tagged EGL-5, which is expressed in PLM, but not ALM (Ferreira *et al.*, 1999; Zheng *et al.*, 2015a) and dissociated cells from these embryos. Based on the expression of EGL-5::GFP, the majority of TRNs *in vitro* appear to adopt an ALM-like cell fate, both in terms of their unipolar morphology and a lack of detectable EGL-5 expression. Sixty-eight percent of TRNs were unipolar; most of these neurons (78%) also did not express EGL-5, consistent with an ALM cell fate (Figure 6A). The remaining cells were bipolar; slightly more than half (62%) of these cells expressed EGL-5, as expected for a PLM cell fate. Given that embryos contain equal numbers of ALM and PLM cells, we performed a chi-squared goodness of fit test to compare our ratios of unipolar:bipolar and EGL-5^+^:EGL-5^−^ to a hypothetical 1:1 ratio. We found that the differences were significantly biased toward ALM morphology and away from EGL-5 expression (*P <*0.001). Thus, embryonic TRNs grown in culture are more likely to adopt an ALM-like cell fate, consistent with the idea that ALM is the default cell fate for TRNs (Zheng *et al.*, 2015a).

### TRN process outgrowth is *pmk-3* p38 MAPK-independent

TRNs generate processes embryonically (Gallegos and Bargmann, 2004; Hilliard and Bargmann, 2006). Similar to *ex vivo* isolation procedures in dissociated spinal ganglia (Daniels, 1972), the embryonic dissociation procedure removes neural processes such that only cell bodies are present in cultures immediately after plating. Because *C. elegans* neurons can regenerate following laser-induced axotomy *in vivo* (Yanik *et al.*, 2005), we reasoned that process outgrowth could recapitulate either a regenerative process or a developmental outgrowth process. As a first step towards distinguishing between these possibilities, we cultured TRNs from animals with a mutation in the p38 MAP kinase *pmk-3,* which is required for process regeneration (Hammarlund *et al.*, 2009; Yan *et al.*, 2009) but not for TRN outgrowth *in vivo* (not shown). The *pmk-3* TRNs extended processes normally *in vitro*; in fact, their processes were slightly longer than wild type cells in the case of the unipolar cells (Figure 6B). Thus, the *in vitro* growth process is independent of *pmk-3* and likely to differ from process regeneration *in vivo*. These results suggest that TRN process growth *in vitro* is more closely related to developmental outgrowth than it is to regeneration.

### TRNs require *mec-7* and *mec-12* for outgrowth *in vitro*

Next, we tested whether or not microtubules were required for outgrowth *in vitro* by treating control cultures with 1 µM colchicine. Although GFP-tagged cell bodies were readily detected in colchicine-treated cultures, we did not detect processes in three independent replicates. In contrast, cultures treated side-by-side with vehicle (DMSO) had processes that were qualitatively and quantitatively similar to untreated controls—the geometric means of their lengths were: unipolar, 41.5 µm; major bipolar, 42.3 µm; minor bipolar, 16.3 µm. Thus, colchicine treatment has stronger effects on TRN process formation *in vitro* than it does *in vivo* (see Figure 1). Next, we prepared cultures from the tubulin mutant strains that we examined in our *in vivo* experiments. Mutations in *tbb-1* and *tbb-2* did not affect the distribution of process lengths. Similar to our observations *in vivo,* however, *mec-7* caused a substantial shift in the distributions of process lengths to shorter distributions. Whereas only PLM was affected *in vivo*, both unipolar and bipolar cells were affected *in vitro*. Process lengths in *tbb-1;mec-7* and *tbb-2;mec-7* double mutants were similar those found in *mec-7* single mutants. In contrast with our *in vivo* observations (Figure 3), we found that loss of *mec-12* caused a strong reduction in the lengths of both unipolar and bipolar cells (Figure 7). Process lengths in *tba-1;mec-12* and *tba-2;mec-12* double mutants were not significantly different from *mec-12* single mutants, however. The differential effect of loss of *mec-12* function *in vivo* and *in vitro* indicates that the roles of the tubulin isoforms crucially depend on their cellular context.

**Figure 7.**
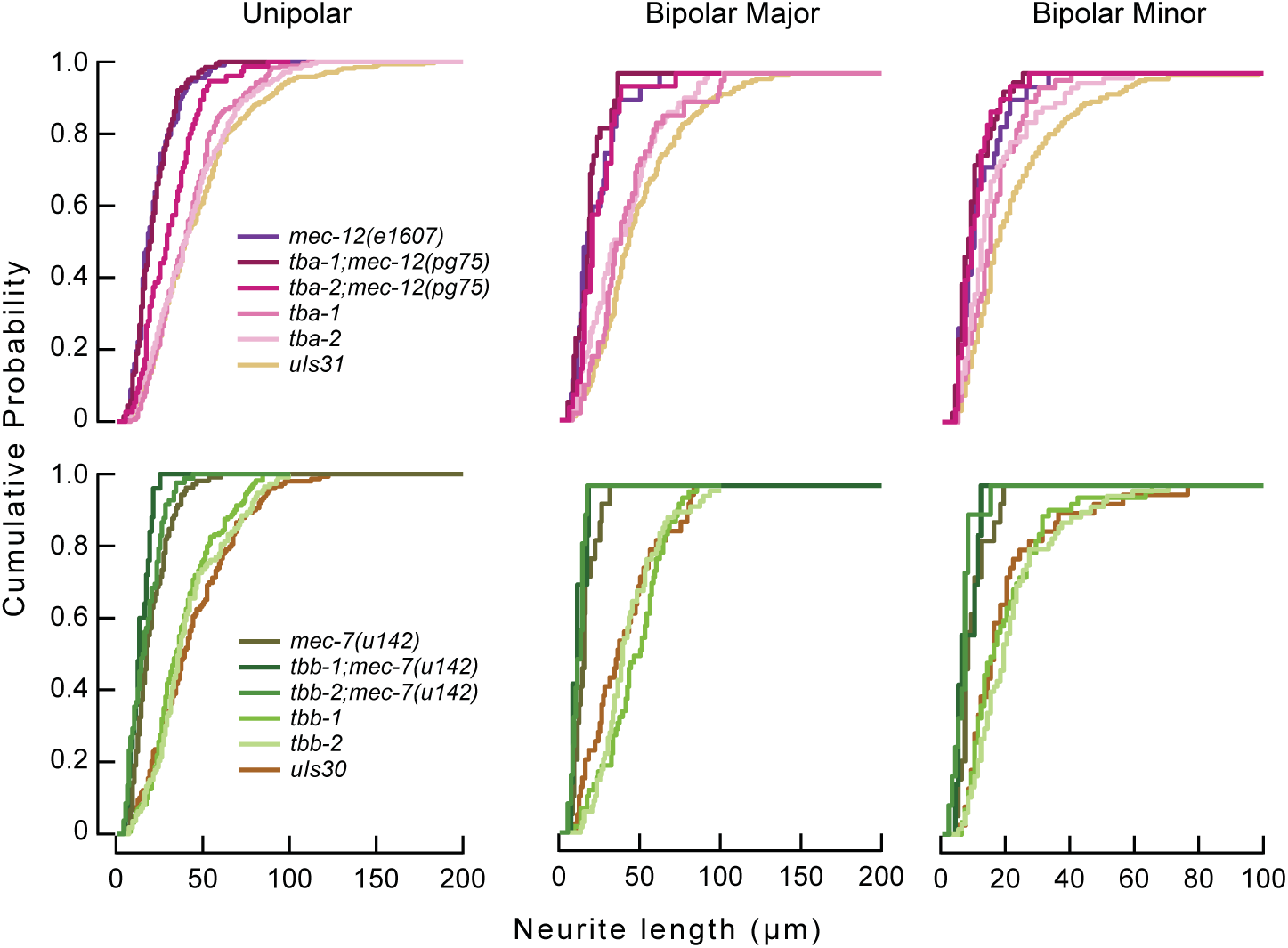
*mec-7* and *mec-12* are necessary for wild-type process outgrowth *in vitro*. Cumulative probability distributions for TRNs isolated from animals of given genotypes plotted by α—tubulin mutants (top row) and β—tubulin mutants (bottom row). Genotypes plotted with GFP controls for majority of corresponding genotypes. Exceptions are *mec-12* and *mec-7*, which are plotted by tubulin family (α or β) for comparisons within class. No statistically significant differences were detected between *uIs30* and *uIs31* GFP controls.

## Discussion

Animal genomes harbor multiple genes encoding α-and β-tubulins that coassemble to form microtubules essential for the growth of long, thin axons and dendrites in the nervous system. Although highly conserved overall, variable portions of α-and β-tubulins diverge within genomes and this finding has fostered the multi-tubulin hypothesis. Our results support a model in which each neuronal type expresses a distinctive palette of tubulins and some tubulin isoforms have evolved for significant redundancy, while others have acquired specialized roles essential for the function of the particular neuron type. Whereas redundancy among tubulin isoforms defends neurons against mutation of single tubulin genes, specialization enables the differentiation of microtubule structures between neuronal subtypes.

For example, the α-tubulins, TBA-1 and TBA-2, are individually dispensable for the outgrowth and function of the touch receptor neurons, while MEC-12 is important for sensory function of TRNs and probably plays a role in neurite outgrowth that is partially compensated by TBA-1 *in vivo*, but revealed *in vitro*. Likewise, the β-tubulins TBB-1 and TBB-2 are individually dispensable for process outgrowth, but loss of MEC-7 causes sensory defects and impairs outgrowth in living animals and shortens neurites in culture. The TRNs are not the only cells that rely on the TBA-1 and TBA-2 α-tubulins and the TBB-1 and TBB-2 β-tubulins. These four tubulins cooperate in a partially redundant manner to enable mitosis and embryogenesis (Phillips *et al.*, 2004). The TBA-1 α-tubulin, along with the TBB-1 and TBB-2 β-tubulins, also contribute to motor neuron axon outgrowth and guidance (Baran *et al.*, 2010). We infer from these results that TBA-1 and TBA-2 α-tubulins and TBB-1 and TBB-2 β-tubulins are incorporated not only into microtubules forming mitotic spindles, but also into microtubules that regulate process outgrowth in neurons.

Because the lateral TRNs assume stereotyped shapes and sizes *in vivo* and arise in the embryo, they provide a fruitful basis for understanding which features of neurite outgrowth are preserved when neurons are dissociated from embryos and grown *in vitro.* For instance, most cultured TRNs have simple bipolar or monopolar shapes similar to those found *in vivo*. But, neurite lengths are far more variable *in vitro* than they are *in vivo*, as evidenced by an approximately 10-fold higher coefficient of variation *in vitro*. More than half of the lateral TRNs adopt an ALM-like, EGL-5-negative cell fate *in vitro*. Additionally, loss of MEC-12 α-tubulin shortens neurites *in vitro*, but not *in vivo*. One important factor that could be linked to all of these features is the presence of native ECM *in vivo* and its absence *in vitro*. For instance, the ECM might compensate for defects in microtubules as seen in colchicine-treated model neurons co-cultured with an ECM-like material (Lamoureux *et al.*, 1990). While cell-cell and cell-matrix signaling is dispensable for outgrowth under many conditions, it could also promote growth, provide a reliable stop-growth signal, and promote expression of *egl-5*.

Tubulins that function redundantly in neurons, as well as compensation from surrounding tissues, could enable the divergence of tubulin genes giving rise to special-purpose tubulins serving neuron-specific functions. In humans, mutations in tubulin genes have been linked to severe defects in brain development or tubulinopathies (Romaniello *et al.*, 2015; Chakraborti *et al.*, 2016). Although most of the currently known tubulinopathies result from missense mutations (Chakraborti *et al.*, 2016), loss of special-purpose tubulins could alter axon guidance and outgrowth, as found for the loss of MEC-7 in *C. elegans* TRNs. Our results suggest that understanding the interplay between cellular context and tubulin isoform function may offer new insights into how neurons regulate a wide array of growth phenotypes from a pool of similar building blocks.

## Methods

### *C. elegans* strains and genetics

We used standard techniques (Brenner, 1974) to maintain strains, which were obtained from the Caenorhabditis Genetics Center (CGC), the Japanese Knockout Consortium (FXO6948), as a generous gift from C, Zheng and M. Chalfie (Columbia University, Department of Biological Sciences; TU4008), or created for the purposes of this study. Table S1 lists the strains used in this study and their provenance. To visualize TRNs *in vivo* and *in vitro*, we used transgenes *uIs31[mec-17p::gfp]III* (O’Hagan *et al.*, 2005) or *uIs30[mec-17p::gfp]I* (Bounoutas *et al.*, 2009) that express GFP selectively in the TRNs. To identify *egl-5*-expressing TRNs in dissociated cell cultures, we used the *uIs115[mec-17p::rfp]* and *uIs116[egl-5p::egl-5::gfp]* transgenes that selectively label TRNs and EGL-5 expressing cells, respectively (Zheng *et al.*, 2015b).

### CRISPR/Cas9-mediated gene disruption

We used CRISPR/Cas9-mediated mutagenesis to create double tubulin knockouts and to combine tubulin gene loss-of-function with transgenes labeling the TRNs with GFP. In particular, we generated *pg75,* a new *mec-12* loss-of-function allele, in the *uIs31* background as follows. (Both *mec-12* and the *uIs31* TRN::GFP transgene are located on chromosome III.) We injected purified *in vitro*-generated CRISPR-Cas9 complexes with sgRNA against the target sequence (5'-GCAGTTTGTCTGCTTTTCCG-3') into the gonads of adult TU2769 *uIs31*[*mec-17p::gfp]* hermaphrodites, essentially as described by Paix et al. (2015). The sgRNA was generated by *in vitro* transcription of a short PCR product containing the target sequence. Recombinant Cas9 was purchased from PNA Bio (#CP01). Individual Mec (Mechanosensory Abnormal) F2 progeny were isolated and their offspring were genotyped to characterize the molecular defect present in the *mec-12* gene. The *pg75* allele corresponds to an 8-nt deletion of TCCGTGGA near the 5’ end of the target site, resulting in a stop codon at amino acid position 317. The *pg75* allele failed to complement the previously identified null allele *mec-12(e1607)* (Fukushige *et al.*, 1999) and was recessive to the wild-type allele in N2. *pg75/e1607* transheterozygotes responded to 1/10 touches (*n=5*) and *pg75/+* heterozygotes responded to 8.5/10 touches (*n=5*). The latter is comparable to the results for the *uIs30* and *uIs31* markers alone (Figure 4). The *pg75* allele was outcrossed twice to minimize the contribution of off-target mutations before using the strain to generate *tba-1(ok1135)I; mec-12(pg75)uIs31[mec-17p::gfp]III* and *tba-2(tm6948)I;mec-12(pg75)uIs31[mec-17p::gfp]III* double mutants.

### CRISPR/Cas9-mediated promoter and protein fusions

To visualize the cells and tissues expressing *tba-1, tba-2,* and *tbb-1*, we adopted the CRISPR/Cas9-mediated genome insertion strategy reported by Dickinson et al. (2013) and obtained GN675 *pg77*[*TagRFP-T*::*tba-1*]; *uIs31*, GN683 *pg79*[*TagRFP-T::tbb-1*]; *uIs31*, and GN688 *pg81*[*TagRFP-T::tba-*2]; *uIs31* (Table S1). (Attempts to insert sequences encoding *TagRFP-T* into the *tbb-2* locus using this strategy were not successful.) For each gene, we generated a sgRNA construct based on the pDD162 plasmid (Addgene #47549) and a homology repair construct that was based on the pDD284 plasmid (Addgene #66825). Briefly, a homology repair construct was generated by PCR with approximately 500 bp of homologous sequences both 5’ and 3’ of the target insertion site flanking the self-excising cassette; this PCR product was then inserted into the pDD284 plasmid using the NEB Hi-Fi Assembly kit (NEB Inc.). We created a Cas9-sgRNA construct by inserting a 20 bp target sequence into the pDD162 plasmid using the Q5 Site-Directed Mutagenesis kit (NEB Inc.). These two constructs were injected into the gonads of young adult hermaphrodites at 10 ng/µL (homology repair) and 50 ng/µL (Cas9 construct). We also injected 50 ng/µL of mg166 (*unc-122p*::RFP), 5 ng/µL of pCFJ104 (*myo-3*p::mCherry::unc-54utr), and 2.5 ng/µL of pCFJ90 (*myo-2p*::mCherry::unc-54utr) as co-injection markers. Putative transgenic animals were cultivated at 25 °C for 3 days; hygromycin was then added to the plates to a final concentration of 250 µg/mL to select for transgenic animals and putative knock-in lines. After 3-5 days of hygromycin selection, we screened for animals that exhibited a strong Rol phenotype, hygromycin resistance, and a lack of co-injection markers as markers for successful insertion into the genome near the PAM site identified for each gene. To generate animals expressing protein fusion constructs, five L1s with the promoter fusion insertions obtained above were picked to each of three NGM plates and subjected to a 34 °C heat shock for 4 hours and then were placed back at 25 °C. After 3 days, we screened for animals that were non-Rol to obtain the fluorescent protein::tubulin fusion animals.

### Tubulin promoter cloning methods

The *tba-1* promoter sequence used in plasmid pRO133 was 1144 base pairs upstream of the genomic *tba-1* start codon. The *tba-2* promoter sequence used in plasmid pRO134 was 1018 base pairs upstream of the genomic *tba-2* start codon. The *tbb-2* promoter sequence used in plasmid pRO137 was 867 base pairs upstream of the genomic *tbb-2* start codon. pRO133, pRO134, and pRO137 were made by fusing promoter sequences to *gfp* coding sequence and *unc-54* 3’ UTR using Gibson Assembly (Gibson *et al.*, 2009).

All tubulin promoter transgenes were made by germline injection transformation of strain PS2172 [*pha-1(e2123) III; him-5(e1490)V*] with an injection mix containing the following DNA: tubulin promoter gfp plasmid at 20ng/µL and *pha-1(+)* rescue plasmid (pBX; Miyabayashi *et al.*, 1999) at 50ng/µL and *unc-122p::rfp* plasmid at 50ng/µL as transformation markers. These transgenics were selected by maintaining animals at ≥ 20**°**C, the restrictive temperature for the *pha-1(e2123)* mutation.

### Single-neuron RNA-seq

Microdissection and single-cell RT-PCR of individual PLM, AFD, and ASER neurons was performed essentially as described (Schwarz *et al.*, 2012). In particular, we dissected and harvested individual GFP^+^ neurons from either L4 or adult animals, separately amplified their transcripts with RT-PCR, and purified and quantitated the RT-PCR products. To determine consensus expression patterns for each neuron type, we pooled aliquots (of varying volume but constant DNA content) from purified and quantitated individual RT-PCR products before performing RNA-seq on the pools. For PLM, we pooled three aliquots; for AFD, seven aliquots, and for ASER, seven aliquots. For all three neuronal types, single-end 50-nt RNA-seq was performed on an Illumina HiSeq 2000. To identify so-called housekeeping genes and genes primarily active outside the nervous system, we compared results from PLM, AFD and ASER to published single-end 38-nt RNA-seq data from mixed-stage whole *C. elegans* hermaphrodite larvae (Schwarz *et al.*, 2012).

Reads were quality-filtered as follows: neuronal reads that failed Chastity filtering were discarded (Chastity filtering had not been available for the larval reads); raw 38-nt larval reads were trimmed 1 nt to 37 nt; all reads were trimmed to remove any indeterminate ("N") residues or residues with a quality score of less than 3; and larval reads that had been trimmed below 37 nt were deleted, as were neuronal reads that had been trimmed below 50 nt. This left a total of 19,316,855 to 27,057,771 filtered reads for analysis of each neuronal type, versus 23,369,056 filtered reads for whole larvae (Table S2).

We used RSEM version 1.2.17 (Li and Dewey, 2011) with bowtie2 version 2.2.3 (Langmead and Salzberg, 2012) and SAMTools version 1.0 (Li *et al.*, 2009) to map filtered reads to a *C. elegans* gene index and generate readcounts and gene expression levels in transcripts per million (TPM). To create the *C. elegans* gene index, we ran RSEM's *rsem-prepare-reference* with the arguments "*--bowtie2 --transcript-to-gene-map*" upon a collection of coding DNA sequences (CDSes) from both protein-coding and non-protein-coding *C. elegans* genes in WormBase release WS245 (Harris *et al.*, 2014). The CDS sequences were obtained from the following sites:

- *ftp://ftp.sanger.ac.uk/pub2/wormbase/releases/WS245/species/c_elegans/PRJNA13758/c_elegans.PRJNA13758.WS245.mRNA_transcripts.fa.gz*.
- *ftp://ftp.sanger.ac.uk/pub2/wormbase/releases/WS245/species/c_elegans/PRJNA13758/c_elegans.PRJNA13758.WS245.ncRNA_transcripts.fa.gz*.

For each RNA-seq data set of interest, we computed mapped reads and expression levels per gene by running RSEM's *rsem-calculate-expression* with the arguments "*--bowtie2 -p 8 --no-bam-output --calc-pme --calc-ci --ci-credibility-level 0.99 --fragment-length-mean 200--fragment-length-sd 20 --estimate-rspd --ci-memory 30000*". These arguments, in particular "*--estimate-rspd*", were aimed at dealing with single-end data from 3'-biased RT-PCR reactions; the arguments "*--phred33-quals*" and "*--phred64-quals*" were also used for the neuronal and larval reads, respectively. We computed posterior mean estimates (PMEs) both for read counts and for gene expression levels, and rounded PMEs of read counts down to the nearest lesser integer. We also computed 99% credibility intervals (CIs) for expression data, so that we could use the minimum value in the 99% CI for TPM as a reliable minimum estimate of a gene's expression (minTPM).

We observed the following overall alignment rates of the reads to the WS245 *C. elegans* gene index: 50.71% for the PML read set, 28.42% for the AFD read set, 31.13% for the ASER read set, and 76.41% for the larval read set (Table S2). A similar discrepancy between lower alignment rates for hand-dissected linker cell RNA-seq reads versus higher alignment rates for whole larval RNA-seq reads was previously observed, and found to be due to a much higher rate of human contaminant RNA sequences in the hand-dissected linker cells (Schwarz *et al.*, 2012). We defined non-zero, above-background expression for a gene in a given RNA-seq data set by that gene having a minimum estimated expression level (termed minTPM) of at least 0.1 TPM in a credibility interval of 99% (i.e., ≥ 0.1 minTPM). The numbers of genes being scored as expressed in a given neuronal type (PLM, AFD, or ASER) above background levels, for various data sets, are given in Table S3. Other results from RSEM analysis are given in Table S4.

We annotated *C. elegans* genes and the encoded gene products in several ways (Table S4). For the products of protein-coding genes, we predicted classical signal and transmembrane sequences with Phobius 1.01 (Käll *et al.*, 2004), regions of low sequence complexity with pseg (SEG for proteins), from *ftp://ftp.ncbi.nlm.nih.gov/pub/seg/pseg* (Wootton, 1994), and coiled-coil domains with ncoils from *http://www.russelllab.org/coils/coils.tar.gz* (Lupas, 1996). PFAM 27.0 protein domains from PFAM-A (Finn *et al.*, 2014) were detected with HMMER 3.0/hmmsearch (Eddy, 2009) at a threshold of E ≤ 10^−5^. The memberships of genes in orthology groups from eggNOG 3.0 (Powell *et al.*, 2012) were extracted from WormBase WS245 with the TableMaker function of ACEDB 4.3.39. Genes with likely housekeeping status (based on ubiquitous expression in both larvae and linker cells) were as identified in our previous work (Schwarz *et al.*, 2012).

### Embryonic cell culture

We isolated cells from *C. elegans* embryos and maintained them in mixed cultures, as described (Strange *et al.*, 2007). Briefly, we treated fifteen 6-cm plates full of synchronized adult animals with sodium hypochlorite hypochlorite (Sulston and Hodgkin, 1988) to obtain an approximately 50 µL pellet of embryos. These embryos were washed with egg buffer (118 mM NaCl, 48 mM KCl, 2 mM CaCl_2_, 2 mM MgCl_2_, 25 mM HEPES, pH 7.3) and separated from debris using a sucrose gradient centrifugation. The embryos were then digested with chitinase for 85 minutes (1 unit, Sigma No. C7809), resuspended in egg buffer (1 mL), triturated (15 times) with a micropipette with the tip pressed against the inside of a microcentrifuge tube, and filtered through a 5-µm syringe filter. The resulting cells were counted with a hemocytometer, resuspended in nematode culture medium (L-15 without Phenol Red, 10% heat-inactivated FBS, 50 U/mL penicillin and 50 mg/mL streptomycin) and plated on glass-bottomed Petri dishes (MatTek). Culture dishes were pre-treated with peanut lectin (0.5 mg/mL) solution for 17 minutes. Cells density was adjusted to 500,000 to 1,000,000 cells/mL and between 50,000 and 100,000 cells were plated on each dish. Cells were maintained in culture at room temperature for 3 days before analysis.

### Imaging TRNs

All of the images aside from the whole worm tubulin expression images (Figure S2) were captured on an inverted Nikon Eclipse Ti-E microscope (Nikon Corporation) equipped with either a PlanApo 100x/1.4 NA oil (*egl-5* imaging), a PlanApo 60x/1.4 NA oil (CRISPR tubulin expression imaging), a PlanApo 40x/0.95 NA air (*in vitro* length measurements), a PlanFluor 40x/1.3 NA oil (*in vitro* length measurements), or a PlanApo 20x/0.75 NA air (*in vivo* length measurements) objective (all objectives manufactured by Nikon). For most *in vivo* imaging, animals were immobilized on 5% agarose pads as reported by Kim et al., (2013), with the exception of the whole worm tubulin expression imaging where the animals were anaesthetized with 10 mM levamisole and mounted on agar pads for imaging at room temperature. Images were captured with a sCMOS camera (Neo 5.5, Andor Technology) controlled by the Micromanager software package (Edelstein et al, 2014) with 4 by 4 pixel binning, or in the case of the whole worm expression imaging, with a Zeiss Axio Imager.D1M (Zeiss, Oberkochen, Germany) with a Retiga-SRV Fast 1394 digital camera (Q-Imaging, Surrey, BC, Canada) or a Zeiss Axioplan2 microscope with a Photometrics Cascade 512B CCD camera, both of which are equipped with 10x, 63x (NA 1.4), and 100x (NA 1.4) oil-immersion objectives.

GFP or RFP expressing TRNs were classified as either unipolar or bipolar prior to analysis. Cells with either one process or with two processes on which the second process was less than one cell body length were classified as unipolar. Cells with two processes that were both greater than one cell body length were classified as bipolar. Cultures also included TRNs with irregular spider web morphologies, but these were rare (<10%) and not analyzed in detail. Process lengths were measured with the segmented line tool in Fiji (Schindelin *et al.*, 2012) after a calibration using a micrograph of a stage micrometer. The scaling conversions were (in pixels/µm): 20x air (0.78); 40x air (1.52); 40x oil (1.54); 60x oil (2.34); 100x oil (3.80). Animals were measured from the opening of the mouth to the point at which the tail narrowed to a thin, hair-like structure due to the difficulty of accurately determining the final termination point of the tail. Thus, our total worm length measurements are systematically smaller than the actual length of the animal from the mouth to the tip of the tail. For cell-fate characterization, we performed *in vitro* cell isolations on TU4008 as described above. TRNs were classified according to their morphology (unipolar or bipolar) and the presence or absence of EGL-5::GFP.

### Drug treatments

Animals were exposed to colchicine throughout development by placing eggs on standard growth plates supplemented with colchicine (1 mM), as described (Chalfie and Thomson, 1982). This concentration was below the 2 mM threshold at which animals become uncoordinated or lethargic (Chalfie and Thomson, 1982). Cultured cells were exposed to colchicine (1µM) via the culture medium. Colchicine was prepared as stock solution in DMSO (40 mM) and its effects were compared to vehicle (DMSO) alone.

### Touch assays

Touch sensitivity was assayed as described (Chalfie *et al.*, 2014). Briefly, adult hermaphrodites (aged one day after their final larval molt) were brushed with an eyelash hair glued to a thin wooden rod. Each animal was tested with 10 stimuli: five to the anterior of the animal and five to the posterior. Responses were considered positive if they elicited either a pause or a reversal. For each genotype tested, a total of 75 bleach-synchronized animals were assayed in cohort of 25 animals across three replicates. All assays were conducted blinded to genotype or drug treatment.

### Statistical methods

Statistical analyses were performed with PRISM (GraphPad Software, La Jolla CA). Process length distributions, both *in vivo* and *in vitro*, were tested for normality using the Shapiro-Wilks test. For *in vivo* analysis of process length, we used the following statistical tests: for colchicine treatment experiments, cell parameters were compared to controls using a t-test. In the case of the *in vivo* TRN imaging experiments or the gentle touch assays, a one-way ANOVA was used to compare the mutant groups to their GFP marker controls. In instances in which the results of the ANOVA rejected the null hypothesis, a post-hoc Dunnett’s test was used to compare each mutant or treatment group against the GFP marker control.

For *in vitro* analysis of neuron morphology and *egl-5* expression, we used to the following statistical tests: comparisons of *in vitro* cell morphology and *egl-5* expression were tested using a chi-squared goodness-of-fit test *vs*. an expected ratio of 50:50 unipolar:bipolar or 50:50 GFP(+):GFP(-), respectively. Distributions of *in vitro* process lengths were compared using the Kolmogorov-Smirnoff test to their GFP controls. Bin sizes for histograms of *in vitro* data were calculated as in (Shimazaki and Shinomoto, 2007) by minimizing the total error between the histogram and the estimated underlying distribution of the data.

### Data Availability

Sequence files for raw RNA-seq reads have been deposited in the NCBI Sequence Read Archive (SRA; *http://www.ncbi.nlm.nih.gov/sra*) under the accession numbers SRR3481679 (PLM neuron pool), SRR3481680 (AFD neuron pool), and SRR3481678 (ASER neuron pool).

## Acknowledgments

This work was funded by: NIH research grants K99NS089942 (to M.K), R01NS047715 (to M.B.G.), R01GM084389 (to P.W.S.), R01DK059418 and R56DK074746 (to M.M.B); The New Jersey Commission on Spinal Cord Research Grant CSCR15IRG014 (to R.O.); and The Stanford Graduate Fellowship and the NIH’s Cellular and Molecular Biology Training Program T32GM007276 (to D.L). P.W.S. is an investigator of the Howard Hughes Medical Institute. We thank the following: Beth Pruitt and Maria Gallegos for discussion and feedback, Zhiwen Liao for numerous microinjections; Chloé Girard for technical consultation; the Japanese Knockout Consortium, Martin Chalfie, and Chaogu Zheng for strains; Igor Antoshechkin at the Jacobs Genomics Laboratory (California Institute of Technology) for RNA-seq; WormBase; and the Goodman, Pruitt, and Dunn groups for insightful discussions regarding this work.

